# GlycanGT: A Foundation Model for Glycan Graphs with Pretrained Representation and Generative Learning

**DOI:** 10.64898/2025.12.14.694171

**Authors:** Akihiro Kitani, Bingyuan Zhang, Koichi Himori, Yusuke Matsui

## Abstract

**Motivation:** Glycans are highly diverse biological sequences, but their functional understanding has lagged behind that of proteins and nucleic acids. Many glycans remain incompletely characterized or ambiguously annotated, limiting computational analyses. Existing computational approaches are primarily graph-based, capturing local structural features but struggling to model global patterns and incomplete sequences.

**Results:** We present GlycanGT, a foundation model for glycans built on a graph transformer architecture. Glycans were represented as graphs with monosaccharides as nodes and glycosidic bonds as edges, and the model was pretrained using a masked language modeling objective. GlycanGT demonstrated higher performance than existing methods across 8 benchmark classification tasks (e.g., 0.734 Macro-F1 in domain prediction and 0.844 AUPRC for immunogenicity classification), and its embeddings formed biologically meaningful clusters that recovered known N- and O-glycan categories. Moreover, GlycanGT accurately proposed candidates for ambiguous sequences, maintaining >80% top-5 accuracy for both monosaccharide and glycosidic bond predictions under high masking levels.

**Availability and implementation:** The pretrained GlycanGT model weights and usage scripts are available on Hugging Face: https://huggingface.co/Akikitani295/GlycanGT. Additional scripts used for analyses in the paper are publicly available on GitHub: https://github.com/matsui-lab/GlycanGT.

## 1 Introduction

Glycans are structurally diverse biomolecules that play essential roles in protein folding, cell–cell communication, and immune regulation (He *et al*., 2024; Varki, 2017; Hao *et al*., 2025; Hart *et al*., 2011; Bektas and Rubenstein, 2011; Gu *et al*., 2012; Wells *et al*., 2001; Pinho *et al*., 2023). Aberrant glycosylation is implicated in many diseases such as cancers (Pinho and Reis, 2015), neurological disorders (Pradeep *et al*., 2023; Zhang *et al*., 2024), and infectious diseases (Zhang and Qu, 2021), yet our understanding of glycans remains far behind that of proteins and nucleic acids due to their nonlinear branching, compositional heterogeneity, and stochastic biosynthesis (Pothukuchi *et al*., 2019). Public repositories such as GlyTouCan (Tiemeyer *et al*., 2017; Fujita *et al*., 2021) contain over 250, 000 entries, but many sequences remain incomplete or ambiguously annotated, underscoring the need for computational methods to capture glycan complexity.

Early computational approaches relied on motif-based features (Li *et al*., 2022; Bao *et al*., 2021; Coff *et al*., 2020; Porter *et al*., 2010), while more recent machine learning methods have leveraged sequence models and graph neural networks (GNNs). RNN-based language models (e.g., SweetTalk (Bojar *et al*., 2021)) treated glycans as “glycowords,” and GNN-based methods (e.g., SweetNet (Burkholz *et al*., 2021), GIFFLAR (Joeres and Bojar, 2024), and GlycanAA (Xu *et al*., 2025)) represented glycans as graphs, achieving improved performance in multiple prediction tasks (e.g., taxonomy, glycosylation, and immunogenicity). However, GNNs suffer from over-smoothing—where node features become indistinguishable as layers deepen—and over-squashing—where long-range dependencies are compressed into limited representations—both of which limit their ability to capture global structural patterns (Oono and Suzuki, 2019). Moreover, no foundation model exists that can jointly learn whole-structure representations and resolve ambiguous glycan sequences.

Here, we present GlycanGT, a foundation model based on a graph transformer (Yun *et al*., 2019; Kim *et al*., 2022; Ying *et al*., 2021; Rampášek *et al*., 2022) architecture pretrained with a masked language modeling objective. Unlike GNNs, graph transformers apply attention mechanisms (Vaswani *et al*., 2017) across entire graphs, enabling learning of both local and global structural relationships. GlycanGT achieves state-of-the-art performance across multiple prediction tasks, uncovers biologically meaningful structural clusters, and accurately proposes candidates for incomplete sequences. Collectively, these findings position GlycanGT as the foundation model for glycans, capable of supporting a broad spectrum of downstream tasks—including classification, functional prediction, and sequence completion—as well as contributing to glycan database expansion and translational applications.

## 2 Materials and Methods

### 2.1 Data sources

We obtained pretraining glycans from GlyCosmos (Yamada *et al*., 2020), which provides access to GlyTouCan (Tiemeyer *et al*., 2017; Fujita *et al*., 2021) entries, as of June 17, 2025 (n = 244, 842). Glycans containing ambiguous symbols (“?”) were excluded. For entries with uncertain linkage notations such as “α1-3/5,” we expanded them into all admissible alternatives (e.g., “α1-3” and “α1-5”), treating each alternative as an independent glycan entry. Downstream benchmark datasets (taxonomy, glycosylation, immunogenicity) were taken from GlycanML (Xu *et al*., 2024). In GlycanML, motif frequencies were precomputed and used to perform k-means clustering (k = 10), with eight clusters designated for training and the remainder split into validation and test sets; we adopted these official splits. To prevent data leakage, any glycans overlapping downstream tasks were removed from pretraining, yielding 83, 739 glycans. The full list of glycans used for pretraining, the entries removed due to ambiguous symbols, and the expanded linkage variants are provided in Supplementary Table S1.

### 2.2 Graph construction

Glycans in IUPAC-condensed notation were converted to graphs using glycowork glycan_to_nxGraph function and extracted head–relation–tail (node–edge–node) triples. Monosaccharides were treated as node tokens and glycosidic bonds as edge tokens. Dictionaries for monosaccharides and glycosidic bonds were constructed based on the GlyTouCan v3.0 notation, after removing ambiguous symbols (“?”, “/”). Stereochemistry and modifications (e.g., L/D forms; “GalNAc3, 4, 6Ac3”, “GalNAc3, 4, 6Me3”) were encoded as distinct tokens. Linkages included canonical forms (e.g., “α1-1”) and allowed variants (“1-1”, “1-P-1”, “α1-1α”). Terminal linkages were excluded from prediction targets because they lack sufficient contextual information for reliable modeling. The graph-construction scripts are publicly available at https://github.com/matsui-lab/GlycanGT.

### 2.3 GlycanGT architecture

Here we adopt a TokenGT (Kim *et al*., 2022) -style pure graph transformer (Yun *et al*., 2019; Ying *et al*., 2021; Rampášek *et al*., 2022) tailored to glycans: every monosaccharide (node) and glycosidic bond (edge) is treated as an independent token, with no graph-specific message passing or locality constraints hard-coded. Building upon TokenGT, we further introduced a masked language modeling (MLM) pretraining objective over both node and edge tokens, enabling self-supervised representation learning tailored to glycans. As shown by Kim et al. (2022), each token embedding is the sum of (i) a linear projection of its content features, (ii) orthogonal random features (ORFs) used as node identifiers—a node *v* receives [*P*_*v*_, *P*_*v*_] and an edge (*u*, *v*) receives [*P*_*u*_, *P*_*v*_]—where the mutual orthogonality of *P*_*v*_ allows the model to infer which nodes and edges are connected through attention inner products, and (iii) a trainable type embedding that distinguishes nodes from edges. A trainable [Graph] token is prepended, and all tokens are projected to model width *d*. The sequence is processed by post-norm Transformer encoder blocks, where each sub-layer (multi-head self-attention or MLP) is followed by LayerNorm and a residual connection. Attention weights (normalized dot products) (Vaswani *et al*., 2017) aggregate information across all tokens so the model can capture both local branch motifs and long-range cross-branch relations; multi-head attention encourages diverse structural dependencies. The [Graph] token serves as the glycan representation for downstream classifiers. We instantiated four scales (ss/small/medium/large) by varying depth, width, and heads (Supplementary Table S2). Kim et al. (2022) showed theoretically that TokenGT with node/type identifiers is at least as expressive as a 2-IGN (and thus 2-WL), exceeding message-passing GNNs in expressiveness; empirically, TokenGT outperformed GNN baselines and effectively captured global dependencies via self-attention. Because it avoids message passing, it does not inherit typical over-smoothing issues. Full attention is *O*((*n* + *m*)^2^) with the number of tokens per glycan, where *n* denotes the number of monosaccharide residues (nodes) and *m* the number of glycosidic bonds (edges); this cost remains practical for typical glycan sizes. For identifier construction, we use fixed, seeded orthogonal random features (ORFs) to ensure reproducibility while avoiding Laplacian-based eigendecomposition. Implementation details and hyperparameters are available in Supplementary Table S2 and our public repository.

### 2.4 Pretraining objective

We used masked language modeling (MLM) over both node (monosaccharide) and edge (linkage) tokens. Masking ratios of 5%, 15%, 25%, 35%, 45%, and 55% were evaluated. The masking ratio for the final model was determined based on the Macro-F1 scores for taxonomy and glycosylation tasks, and AUPRC for the immunogenicity prediction tasks (Supplementary Figure S3). The loss was cross-entropy with an edge loss weight of 0.5. Optimization used AdamW (learning rate 1×10⁻⁶, weight decay 0.01). A 90/10 train/validation split with early stopping (no mean validation-loss improvement for 15 epochs for ss/small or 30 epochs for medium/large) determined the number of epochs, after which we retrained on the full set.

#### Ablation

To evaluate the contribution of each token type to model learning, we compared a balanced 35% node–35% edge masking strategy with two single-modality variants: (i) 70% node masking with 0% edge masking, and (ii) 70% edge masking with 0% node masking, ensuring a comparable total masking ratio (∼35%) across settings. These experiments were designed to assess whether jointly masking both token types improves contextual learning. Final-layer attention weights were averaged across heads and visualized as token–token attention matrices to examine modality-specific focus patterns (Supplementary Figure S6d–f).

### 2.5 Downstream task evaluation

For each downstream task, glycans were encoded by the pretrained model and the [Graph] token embedding was used as the feature vector. We adopted three benchmark tasks— taxonomy, glycosylation, and immunogenicity—from the GlycanML dataset (Xu et al., 2024), which provides standardized task definitions and dataset splits. These tasks were chosen because they represent biologically meaningful and well-established prediction problems in glycoinformatics. The taxonomy task involves predicting glycan origin at eight hierarchical levels (domain, kingdom, phylum, class, order, family, genus, and species), comprising 13, 209 glycans from SugarBase with 4–1, 737 classes per level. The glycosylation task aims to classify glycans as N-linked, O-linked, or free, using 1, 683 glycans curated from GlyConnect. The immunogenicity task involves binary classification of 1, 320 glycans annotated for immune activity based on literature evidence in SugarBase. All three tasks employ motif-based dataset splits (8:1:1 for training, validation, and test) to evaluate generalization across structurally distinct glycans. Tasks and metrics were: taxonomy (domain to species; Macro-F1, accuracy), glycosylation (N-linked, O-linked, free; Macro-F1, accuracy), and immunogenicity (binary; AUPRC, accuracy). We trained support vector machines (SVM) (Boser *et al*., 1992) and LightGBM (Ke *et al*., 2017) classifiers. These classifiers were chosen as lightweight yet strong baseline models that are well-established for tabular and structured data. Our primary goal was to evaluate the quality of GlycanGT representations rather than to optimize classifier architectures; thus, we used SVM and LightGBM for their simplicity, robustness, and reproducible performance across diverse datasets. For SVM, we performed randomized hyperparameter search over 10 iterations, testing radial basis function (RBF) and linear kernels with regularization parameter C ∼ log U (10⁻³, 10²) and γ ∼ log U (10⁻⁴, 10⁻¹) for RBF. For LightGBM, we ran 15 randomized search iterations with the following ranges: n_estimators ∈ [100, 1000], learning_rate ∼ log U (0.01, 0.3), num_leaves ∈ [20, 150], max_depth ∈ [5, 20], reg_alpha, reg_lambda ∼ log U (10⁻², 10¹), and colsample_bytree, subsample ∼ U (0.6, 1.0). All classifiers used class-balanced weighting and were trained with consistent random seeds for comparability. We followed the official fixed train/validation/test splits of GlycanML. Within the combined training set (train ∪ validation), hyperparameters were selected by RandomizedSearch with 3-fold cross-validation. The best model was then evaluated once on the held-out test split. To account for stochasticity, we repeated the entire procedure with three different random seeds and reported mean ± s.d. over the three runs, following the evaluation protocol of GlycanML.

### 2.6 Baseline methods

We compared GlycanGT against three representative graph-based models: SweetNet (Burkholz *et al*., 2021), GlycanAA (Xu *et al*., 2025), and an RGCN implementation provided by GlycanML (Xu *et al*., 2024). SweetNet applies GNNs to glycan graphs, focusing on local branching motifs. GlycanAA represents glycans as heterogeneous graphs with atom and monosaccharide nodes and hierarchical message passing between these levels using relational graph convolution (RGConv) (Schlichtkrull *et al*., 2017), enabling multi-scale feature learning. RGCN models multi-relational edges to capture diverse linkage types; we used the official GlycanML implementation (accessed in July 2025) with default hyperparameters as described in Xu et al. (2024). All baselines were trained and evaluated on the same datasets and data splits described in Section 2.5 to ensure fair comparison.

### 2.7 Structural similarity analysis using graph kernels

To evaluate whether GlycanGT captures global structural patterns beyond local substructure similarities, we computed pairwise graph similarities using the Weisfeiler–Lehman (WL) subtree kernel (Nguyen *et al*., 2021) implemented in the GraKeL library. Glycans were first converted from IUPAC-condensed notation into NetworkX graphs using the glycan_to_nxGraph function of glycowork. Each node was labeled by its monosaccharide identity, and edges represented glycosidic linkages. The WL subtree kernel iteratively compares relabeled node neighborhoods to quantify structural similarity across branching patterns. A normalized similarity matrix *K*_global_ was computed for all pairwise glycan combinations. The resulting similarity matrices were visualized as hierarchical cluster heatmaps.

### 2.8 Visualization and motif enrichment analysis

To visualize the structural organization of glycan representations learned by GlycanGT, we projected the pretrained [Graph] token embeddings into two dimensions using t-distributed stochastic neighbor embedding (t-SNE) implemented in scikit-learn. Each glycan was represented by the [Graph] embedding obtained from the pretrained large model without additional fine-tuning. The t-SNE algorithm was initialized by PCA and optimized for 1, 000 iterations with a perplexity of 30. Subsequently, to examine functional relationships among embedding clusters, we conducted motif enrichment analysis. GlycanGT embeddings for the glycosylation dataset were reduced to 50 dimensions by PCA and clustered by k-means (k = 6). Cluster-wise motif enrichment was assessed using glycowork’s get_pvals_motifs function (Thomès *et al*., 2021). For each cluster, IUPAC-condensed glycan names were labeled as target (1) or non-target (0), and motif occurrence ratios were compared using Welch’s t-test with FDR correction and Cohen’s d effect sizes. Motifs with p < 0.05 after FDR correction were considered significantly enriched.

### 2.9 Ambiguous glycan analysis

Because masked language modeling (MLM) enables masked token prediction, we evaluated GlycanGT’s ability to reconstruct incomplete or uncertain glycan sequences resembling real-world ambiguity patterns observed in GlyTouCan (Tiemeyer *et al*., 2017; Fujita *et al*., 2021). We considered two representative scenarios derived from empirical distributions of ambiguous annotations (Supplementary Figure S7b): (i) 0% masking of linkages with 10– 90% masking of monosaccharides, and (ii) 100% masking of linkages with 0–90% masking of monosaccharides. To ensure strict separation from pretraining, we used 5678 glycans excluded from the pretraining set due to compositional overlap with downstream datasets. Predictions were ranked by softmax probabilities over the masked token vocabulary, and performance was measured by hit@K (K = {1, 2, 3, 5, 10, 20, 30}), defined as the fraction of cases where the correct token (monosaccharide or linkage) appeared within the top-K predictions. This experiment was designed to assess the model’s capability for automated reconstruction of incompletely annotated glycans.

### 2.10 Statistics

Unless noted, error bars denote mean ± s.d. across replicates. Model metrics (Macro-F1, AUPRC, accuracy) were averaged over three runs. Multiple testing was corrected by Benjamini–Hochberg.

### 2.11 Reproducibility and computing environment

Experiments ran on NVIDIA L40 GPUs (46 GB memory). Software: Python 3.12, PyTorch 2.5.1 (CUDA 12.1). Codes are provided at https://github.com/matsui-lab/GlycanGT and the pretrained model weights are available on Hugging Face: https://huggingface.co/Akikitani295/GlycanGT.

## 3 Results

### 3.1 Pretraining of GlycanGT using masked language modeling

For model pretraining, glycans with complete structural information were curated from GlyCosmos (Fujita *et al*., 2021; Yamada *et al*., 2020). Glycans containing ambiguous symbols (e.g., “?”) as well as those included in downstream analyses were excluded to avoid data leakage, resulting in a pretraining dataset of 83, 739 glycans. Each glycan was converted into a graph representation, with monosaccharides treated as nodes and glycosidic bonds as edges, and provided as input to TokenGT (Kim *et al*., 2022), a variant of the graph transformer architecture (Yun *et al*., 2019; Ying *et al*., 2021; Rampášek *et al*., 2022) (Figure 1 and Supplementary Figure S1, see Materials and Methods for details).

**Figure 1.**
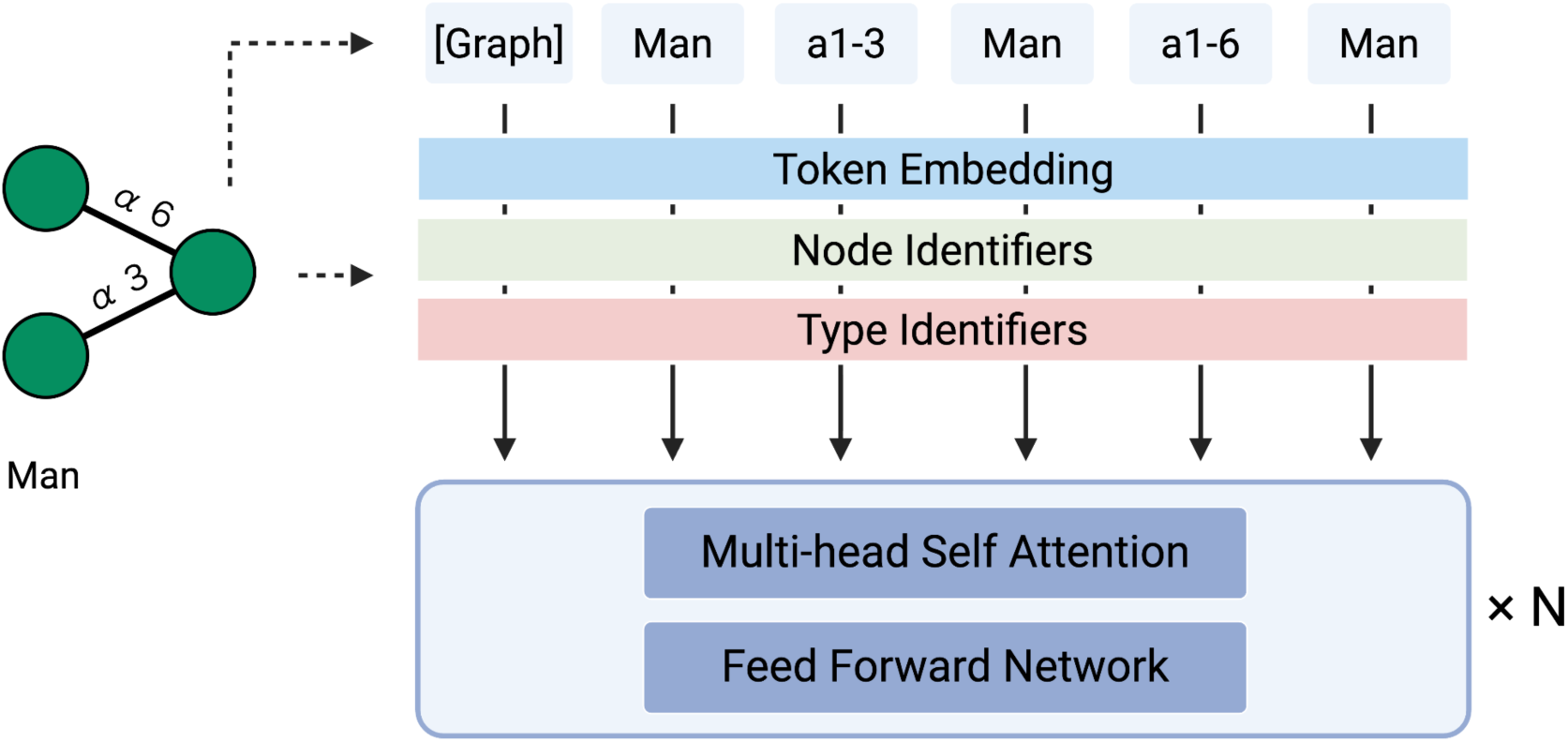
Overview of GlycanGT architecture. Glycans are represented as graphs, where monosaccharides correspond to node tokens and glycosidic bonds to edge tokens. These tokens are processed by a Tokenized Graph Transformer (TokenGT). Each token embedding is constructed as the sum of three components: (1) Token embeddings, derived from token IDs; (2) Node identifiers, orthonormal vectors assigned to nodes to encode graph connectivity; and (3) Type identifiers, trainable vectors indicating whether a token corresponds to a node or an edge. The resulting augmented embeddings are input to the Transformer encoder layers to learn glycan graph representations.

Pretraining was performed using a masked language modeling (MLM) objective, originally developed for natural language processing (Devlin *et al*., 2018), in which monosaccharides and glycosidic bonds were randomly masked at specified probabilities and predicted from the surrounding context. Six masking ratios (5%, 15%, 25%, 35%, 45%, and 55%) were examined (Supplementary Figure S2, Supplementary Table S2 and S3). Four model scales (ss, small, medium, and large) were trained, among which the large model consistently achieved the lowest loss and was therefore used for further exploration of optimal masking ratios.

Embeddings obtained from the pretrained models were applied as input features for downstream prediction tasks, with support vector machines (SVM) (Boser *et al*., 1992) and LightGBM (Ke *et al*., 2017) used for classification. Three types of classification tasks were evaluated: taxonomy, glycosylation, and immunogenicity, using previously published benchmark datasets (Xu *et al*., 2024). Across tasks, masking ratios of 25–45% generally yielded the best performance, with a masking ratio of 35% providing consistently strong results (Supplementary Figure S3). Therefore, we adopted the large model pretrained with a 35% masking ratio for subsequent evaluations.

### 3.2 Benchmarking GlycanGT against existing models

We next evaluated the performance of GlycanGT against previously reported methods (Xu *et al*., 2024). As baselines, we considered SweetNet (Burkholz *et al*., 2021), GlycanAA (Xu *et al*., 2025), and the relational graph convolutional network (RGCN) implemented in GlycanML (Xu *et al*., 2024). For taxonomy classification spanning domain to species, GlycanGT achieved the best Macro-F1 performance at all levels except the class level (Figure 2a). However, when evaluated by accuracy (Supplementary Figure S4), GlycanAA performed best at five taxonomic levels, whereas GlycanGT was best at three levels (see Supplementary Table S4 for details). For glycosylation prediction, GlycanGT yielded a Macro-F1 of 0.932, which was lower than that of GlycanAA (0.952) and RGCN (0.936) (Figure 2b). For immunogenicity prediction, GlycanGT achieved the best performance, with an AUPRC of 0.844, compared with 0.705 for GlycanAA and 0.695 for RGCN. Similar trends were also observed when evaluated using accuracy (Supplementary Figure S4 and Supplementary Table S4).

**Figure 2.**
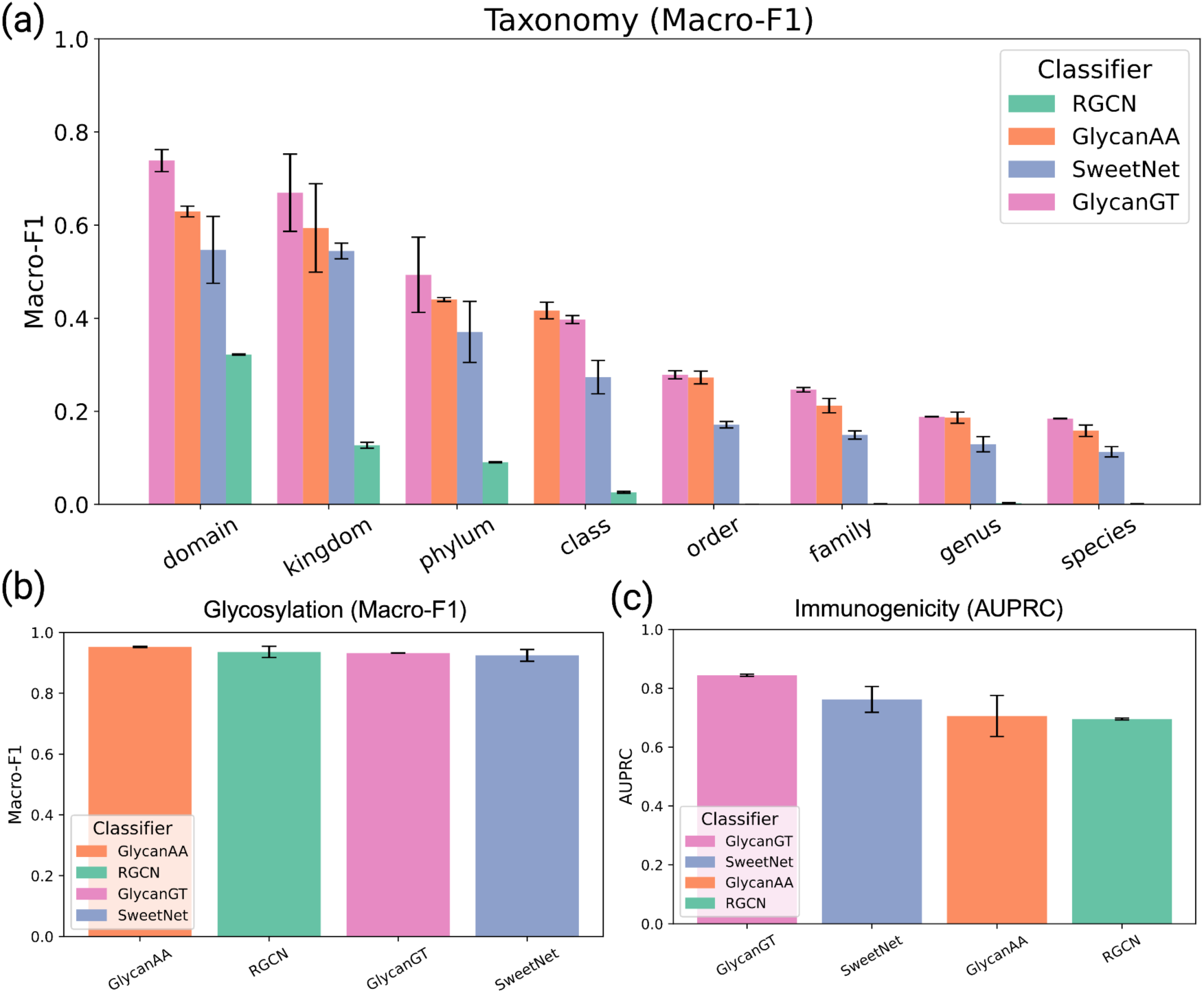
Performance comparison of GlycanGT and baseline models on prediction tasks. (a) Taxonomy classification evaluated by Macro-F1 across eight hierarchical levels (domain to species). (b) Glycosylation prediction evaluated by Macro-F1. (c) Immunogenicity prediction evaluated by AUPRC. Error bars represent mean ± s.d. across replicates.

### 3.3 Global structural patterns captured by GlycanGT

GlycanGT showed high performance in taxonomy classification. We hypothesized that this improvement stems from GlycanGT’s ability to capture the global structural patterns of glycans more effectively than existing GNN-based models. To test this hypothesis, we applied a graph kernel approach to compute pairwise similarities between glycans and examined structural clustering using hierarchical clustering. Specifically, we employed the Weisfeiler–Lehman (WL) subtree kernel (Nguyen *et al*., 2021), which quantitatively compares branching patterns of glycans by evaluating the similarity of their substructures. Using this approach, structural clustering did not recapitulate distinctions based on taxonomy (Supplementary Figure S5a). In contrast, glycans were clustered by glycosylation type (N-linked, O-linked, or free glycans) and by immunogenicity status (immunogenic vs. non-immunogenic) (Supplementary Figure S5b and c). Taken together, these results suggest that taxonomy classification is not strongly dependent on substructural similarities, but rather benefits from the ability of the graph transformer pretraining to capture global structural patterns of glycans.

### 3.4 Biological interpretation via motif enrichment analysis

To assess the effect of pretraining, glycan representations were projected into two dimensions using t-distributed stochastic neighbor embedding (t-SNE) and plotted for each dataset—taxonomy (Figure 3a), glycosylation (Figure 3b), and immunogenicity (Figure 3c). The embeddings effectively separated classes in taxonomy, glycosylation, and immunogenicity tasks, with additional subclusters observed within each class. To further examine their biological relevance, we clustered the glycosylation embeddings and performed motif enrichment analysis, which identified six clusters characterized by distinct glycan motifs (Figure 3d, Supplementary Table S5). Notably, these clusters corresponded to known biological categories. For example, Cluster 0 was enriched for sialylated N-glycan motifs such as NeuNAc(a2-3/6/?)Gal, which are known to promote tumor cell survival, invasion, and immune evasion through enhanced α2-3/6-linked sialylation and interaction with Siglec receptors in the tumor microenvironment (van Vliet and van Kooyk, 2025). In contrast, Cluster 2 contained O-glycan cores initiated by GalNAc such as GlcNAc(b1-6)GalNAc, Gal(b1-3)GalNAc—structures typical of mucin-type glycoproteins involved in epithelial protection and cell–cell adhesion (Bektas and Rubenstein, 2011). Other clusters also captured biologically relevant motifs such as high-mannose, and Plant- and insect-type xylosylated N-glycans. These observations indicate that GlycanGT embeddings preserve functional hierarchies of glycan biosynthesis and organization, capturing biologically meaningful structural diversity that aligns with known N- and O-glycan classes.

**Figure 3.**
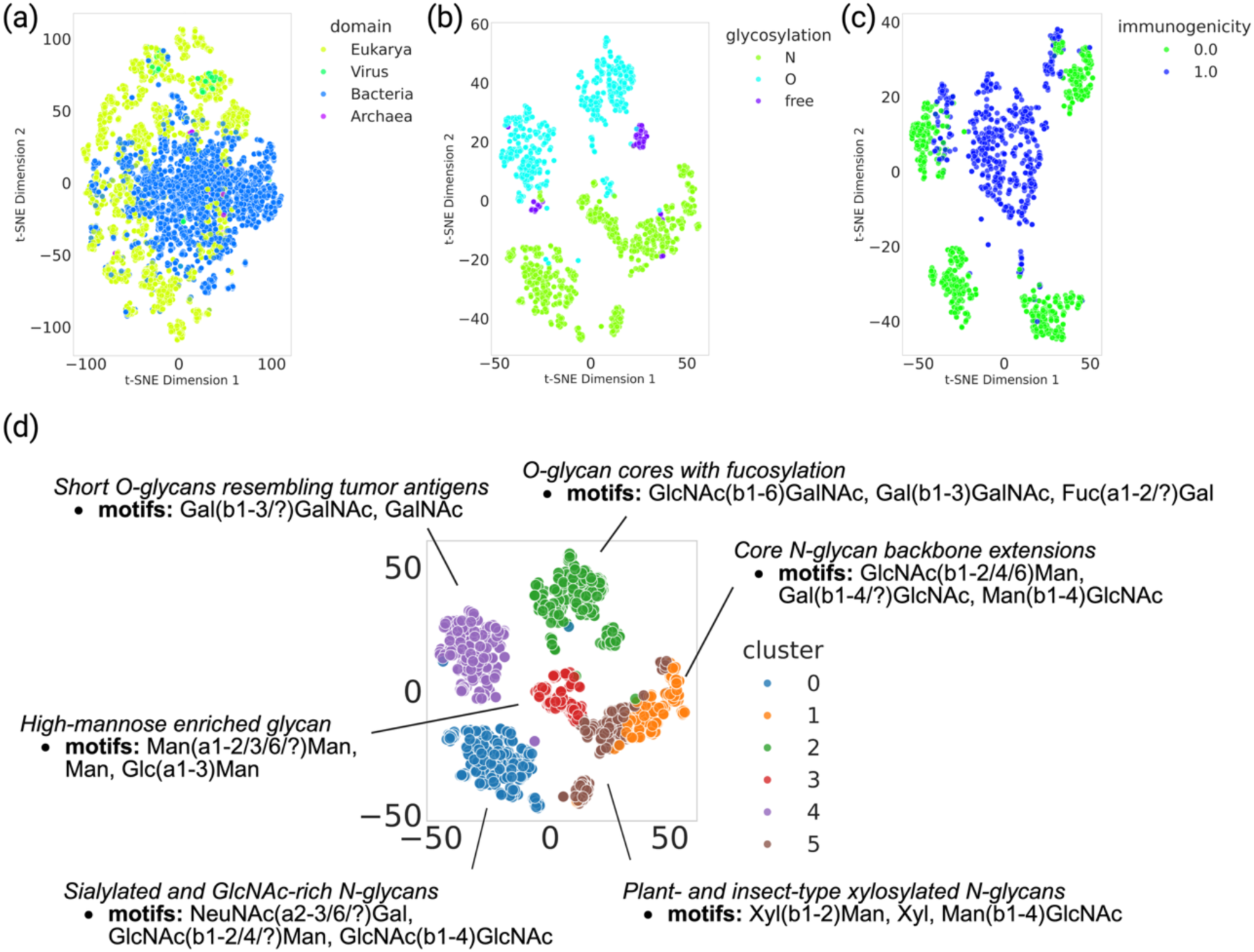
Visualization of glycan representations learned by GlycanGT. Two-dimensional plots of glycan embeddings projected by t-distributed stochastic neighbor embedding (t-SNE). (a) Taxonomy classification at the domain level (eukarya, virus, bacteria, archaea). (b) Glycosylation types (N-linked, O-linked, and free glycans). (c) Immunogenicity (0.0: non-immunogenic, 1.0: immunogenic). (d) Motif enrichment analysis of clusters derived from glycosylation embeddings.

### 3.5 Ablation analysis of masking strategies

To assess the necessity of masking both token types, we compared the balanced 35%–35% masking strategy with two alternatives: 70% monosaccharide-only masking and 70% glycosidic bond-only masking. The balanced strategy consistently yielded superior downstream performance (Supplementary Figure S6a–c). Attention weight visualizations further showed that the balanced model distributed attention across both token types, whereas the single-modality models placed disproportionate weight on the unmasked tokens (Supplementary Figure S6d–f).

### 3.6 Prediction of ambiguous glycan sequences

Many glycans in GlyTouCan contain ambiguous sequence information (e.g., “?” or “/”). As of June 2025, more than half of the registered glycans lacked a defined IUPAC-condensed name, and among those with names, ∼60% contained ambiguous symbols (Supplementary Figure S7a). The distribution of such ambiguities followed two common patterns: (i) unmasked glycosidic bonds with 10–90% of monosaccharides masked, and (ii) fully masked glycosidic bonds with 0–90% of monosaccharides masked (Supplementary Figure S7b).

To evaluate GlycanGT under these conditions, we masked complete glycan sequences excluded from pretraining and assessed prediction accuracy. In the first scenario, top-1 accuracy for monosaccharide prediction remained above 80% across most masking ratios (Figure 4a). In the second, accuracy declined at high masking levels (dropping to ∼50% at 90% masking), but the correct monosaccharide or glycosidic bond appeared within the top-5 predictions in >80% of cases (Figure 4b, c; Supplementary Table S6). These results demonstrate that GlycanGT can propose plausible candidates for ambiguous glycans. Importantly, this capability suggests that GlycanGT could support the curation of glycan databases by filling in missing or uncertain structural elements, as exemplified by our model-based predictions for GlyTouCan entries with ambiguous annotations (Supplementary Table S7).

**Figure 4.**
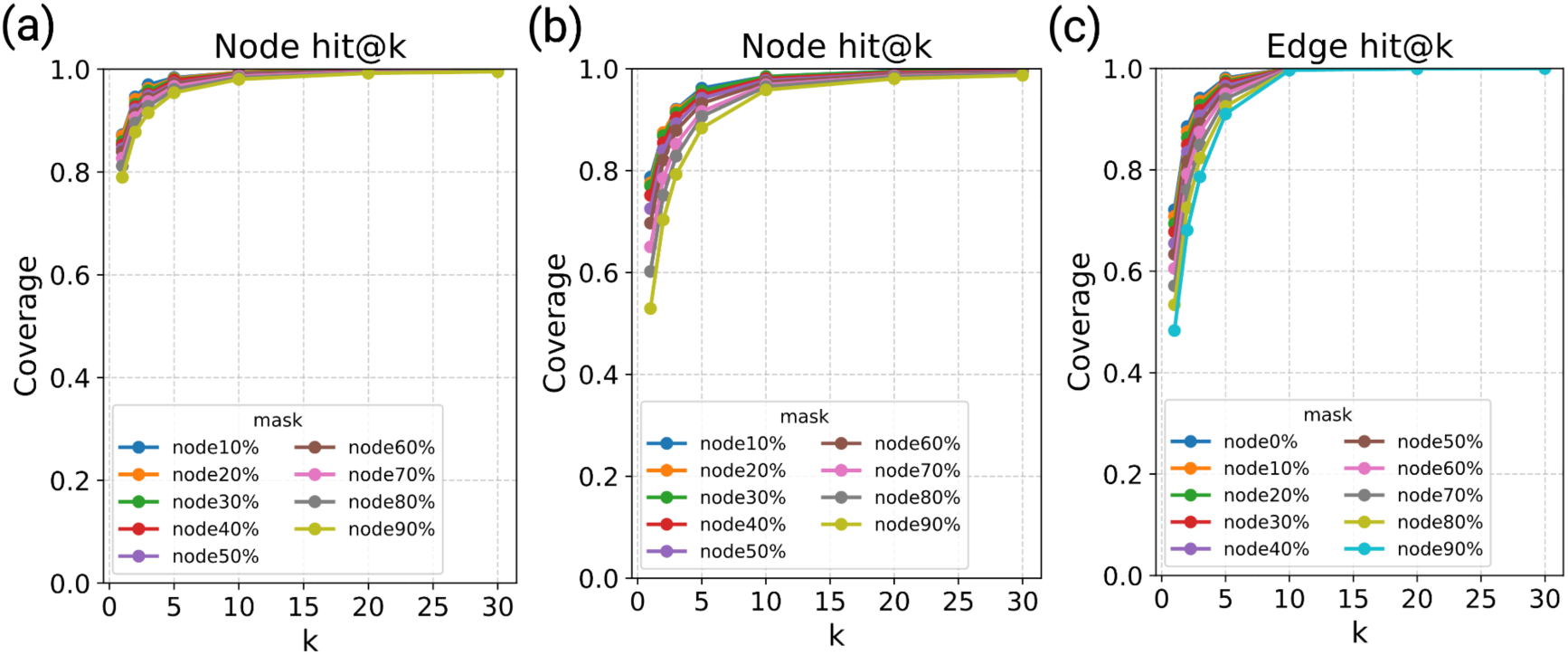
Prediction performance of GlycanGT for ambiguous glycans. (a) Monosaccharide prediction accuracy (hit@K) when glycosidic bonds were not masked (0% masking) and monosaccharide masking ratios varied from 0–90%. (b, c) Prediction accuracy when glycosidic bonds were fully masked (100%). (b) Monosaccharide prediction with varying monosaccharide masking ratios (0–90%). (c) Glycosidic bonds prediction with varying monosaccharide masking ratios (0–90%).

## 4 Discussion

In this study, we developed GlycanGT, a glycan foundation model based on the graph transformer (Yun *et al*., 2019; Kim *et al*., 2022; Ying *et al*., 2021; Rampášek *et al*., 2022) architecture. By employing masked language modeling for pretraining, GlycanGT outperformed previous approaches such as SweetNet (Burkholz *et al*., 2021), GlycanAA (Xu *et al*., 2025), and RGCN across multiple benchmark tasks. Although GlycanAA achieved slightly better performance in certain taxonomy and glycosylation classification tasks, this may reflect its atom- and residue-level feature representations that emphasize local structural motifs. In contrast, GlycanGT focuses on capturing long-range contextual dependencies across entire glycan structures, which likely accounts for its superior performance in taxonomy and immunogenicity prediction and ambiguous sequence completion. Notably, GlycanGT effectively mitigated limitations of conventional GNN-based models, such as over-smoothing and over-squashing (Oono and Suzuki, 2019), by leveraging full self-attention across all node–edge tokens. This architecture removes the locality constraints inherent to message-passing networks and enables efficient propagation of information across distant branches. Similar graph transformer architectures have been theoretically shown to overcome these limitations (Kim et al., 2022; Rampášek et al., 2022), and our benchmarking and structural analyses (Sections 3.2 and 3.3) empirically support this improvement, demonstrating enhanced capture of global structural dependencies and superior predictive performance compared with GNN-based models such as SweetNet and GlycanAA. Furthermore, because it was pretrained with a masked language modeling (MLM) objective, GlycanGT can infer missing structural elements by predicting masked monosaccharides and linkages from their surrounding context. This capability enables automated reconstruction of incompletely annotated glycans, as demonstrated by its >80% Top-1 accuracy and >80% Top-5 hit rates in ambiguous sequence prediction (Section 3.6). Such context-aware completion could facilitate the curation of public databases GlyTouCan (Tiemeyer et al., 2017; Fujita et al., 2021) by suggesting plausible candidates for entries with uncertain or partially defined structures.

Despite these strengths, our study has certain limitations. First, the number of fully defined glycan structures remains limited, and the pretraining dataset used in this work comprised only ∼80, 000 glycans, which may restrict the model’s generalization to rare or synthetic structures. In transformer-based models, performance is known to follow scaling laws with respect to model size, data quantity, and computational budget (Kaplan et al., 2020). Therefore, we expect that pretraining on larger and more diverse glycan datasets will further improve GlycanGT’s performance and generalizability. Second, inconsistencies in glycan notation required stereochemistry and modifications (e.g., L/D forms, acetylation, methylation) as well as linkage variants (e.g., “α1-1”, “1-1”, “1-P-1”) to be treated as distinct tokens. This redundancy can lead to sparse or overlapping representations, potentially affecting embedding stability, even though these variants may represent nearly identical entities. In addition, the quadratic complexity of full self-attention poses computational challenges for scaling to extremely large glycan graphs. These limitations are expected to be mitigated as larger and more standardized glycan datasets become available, and future work will explore scalable attention mechanisms and cross-modal pretraining that integrates glycomics and glycoproteomics data to further enhance model robustness and generalizability.

Overall, GlycanGT demonstrated strong performance across classification tasks and ambiguous sequence completion tasks, achieving up to 0.844 AUPRC in immunogenicity prediction and maintaining >80% Top-1 accuracy for masked monosaccharide prediction (Sections 3.2 and 3.6). The model further provided biologically interpretable embeddings that facilitated motif discovery and immunogenicity analysis. As both pretrained weights and code are publicly available, GlycanGT can be readily fine-tuned or adapted to additional glycan-related datasets and downstream tasks. We envision GlycanGT serving as a reusable foundation model for glycans—bridging structural, functional, and omics-level analyses—and contributing to the mechanistic understanding of disease and the development of glycan-based therapeutic strategies.

## Author contributions

Kitani Akihiro (Conceptualization [lead], Data curation [lead], Formal analysis [lead], Investigation [lead], Methodology [lead], Software [lead], Validation [lead], Visualization [lead], Writing—original draft [lead], Writing—review & editing [lead]), Matsui Yusuke (Supervision [lead], Project administration [lead], Funding acquisition [lead], Writing—review & editing [supporting]), Zhang Bingyuan, Koichi Himori (Conceptualization [supporting], Methodology [supporting], Formal analysis [supporting], Investigation [supporting], Writing— review & editing [supporting])

## Supplementary data

Document S1. Supplementary Figures S1–S8 – pdf file

Supplementary Table S1. Pretraining glycan dataset construction: original IDs, ambiguity removal, and linkage expansion – xlsx file

Supplementary Table S2. Model architectures and number of trainable parameters – xlsx file

Supplementary Table S3. Best validation loss and epochs for GlycanGT models at different masking ratios – xlsx file

Supplementary Table S4. Comparison of GlycanGT with baseline methods – xlsx file

Supplementary Table S5. Motif enrichment analysis of clusters derived from glycosylation embeddings – xlsx file

Supplementary Table S6. Evaluation of GlycanGT prediction performance for ambiguous glycans – xlsx file

Supplementary Table S7. Prediction of ambiguous glycan sequences in GlyTouCan using GlycanGT – xlsx file

Conflict of interest: None declared.

## Data Availability

All data used in this study were obtained from publicly available sources. The pretrained GlycanGT model weights and usage scripts are available on Hugging Face: https://huggingface.co/Akikitani295/GlycanGT. Additional source code for constructing GlycanGT and reproducing the analyses in this study is provided at: https://github.com/matsui-lab/GlycanGT.

