## Supplementary Information for "GlycanGT: A Foundation Model for Glycan Graphs with Pretrained Representation and Generative Learning"

Yusuke Matsui

### Supplementary Figures

### Supplementary Figure S1: Overview of the study

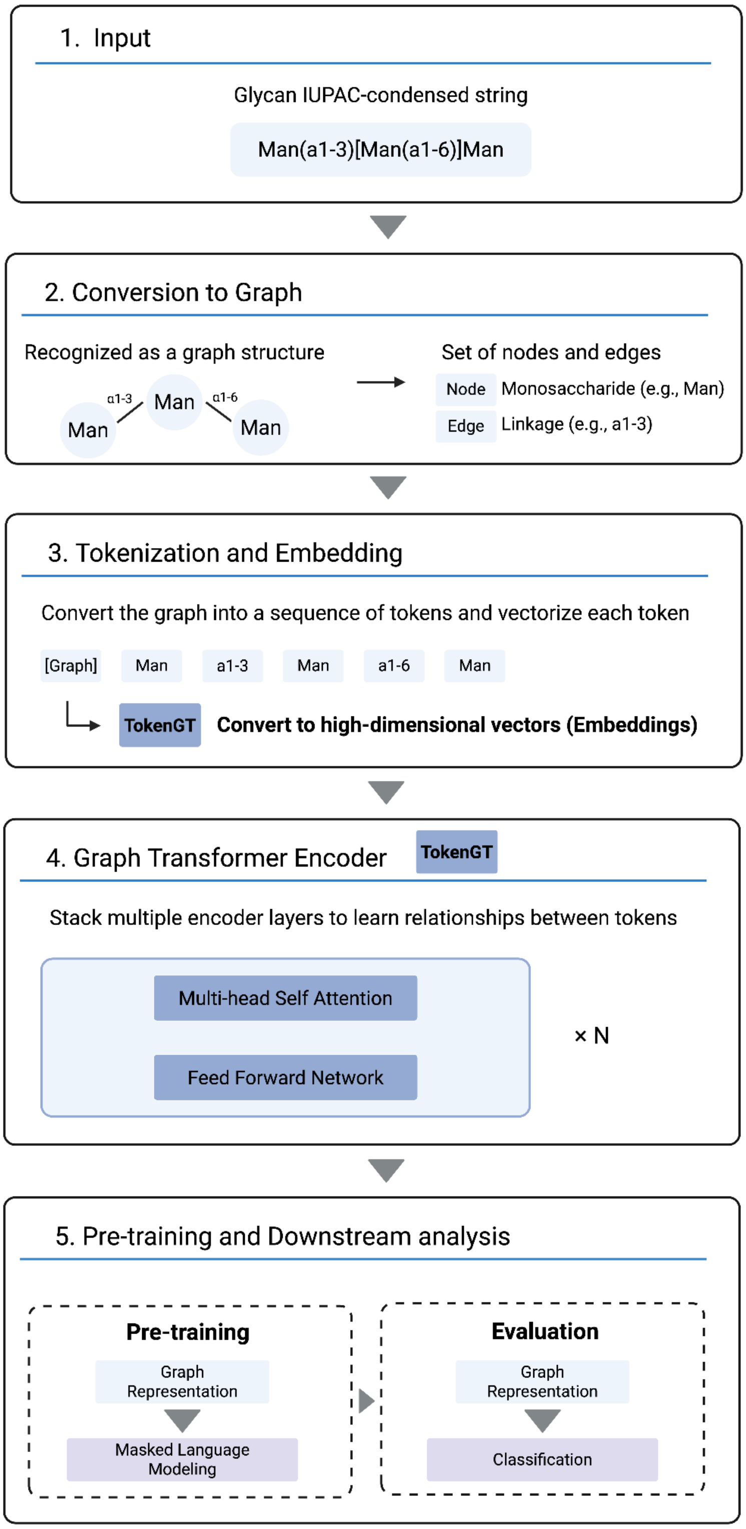

The study consists of five major steps. (1) Input: Pretraining data were obtained from GlyCosmos and represented in IUPAC condensed format. (2) Conversion to graph: Each glycan was converted into a graph where monosaccharides are represented as nodes and glycosidic bonds as edges. (3) Tokenization and embedding: The graph was tokenized using a Graph Transformer architecture (TokenGT) (Kim et al., 2022), and each token was vectorized into a high-dimensional embedding. (4) Graph Transformer encoder: The token embeddings were fed into stacked Transformer encoder layers to learn relationships between tokens. (5) Downstream analysis: The model was pretrained with a masked language modeling objective and evaluated on classification tasks including taxonomy, glycosylation, and immunogenicity prediction.

Supplementary Figure S2: Training and validation loss curves under different masking ratios

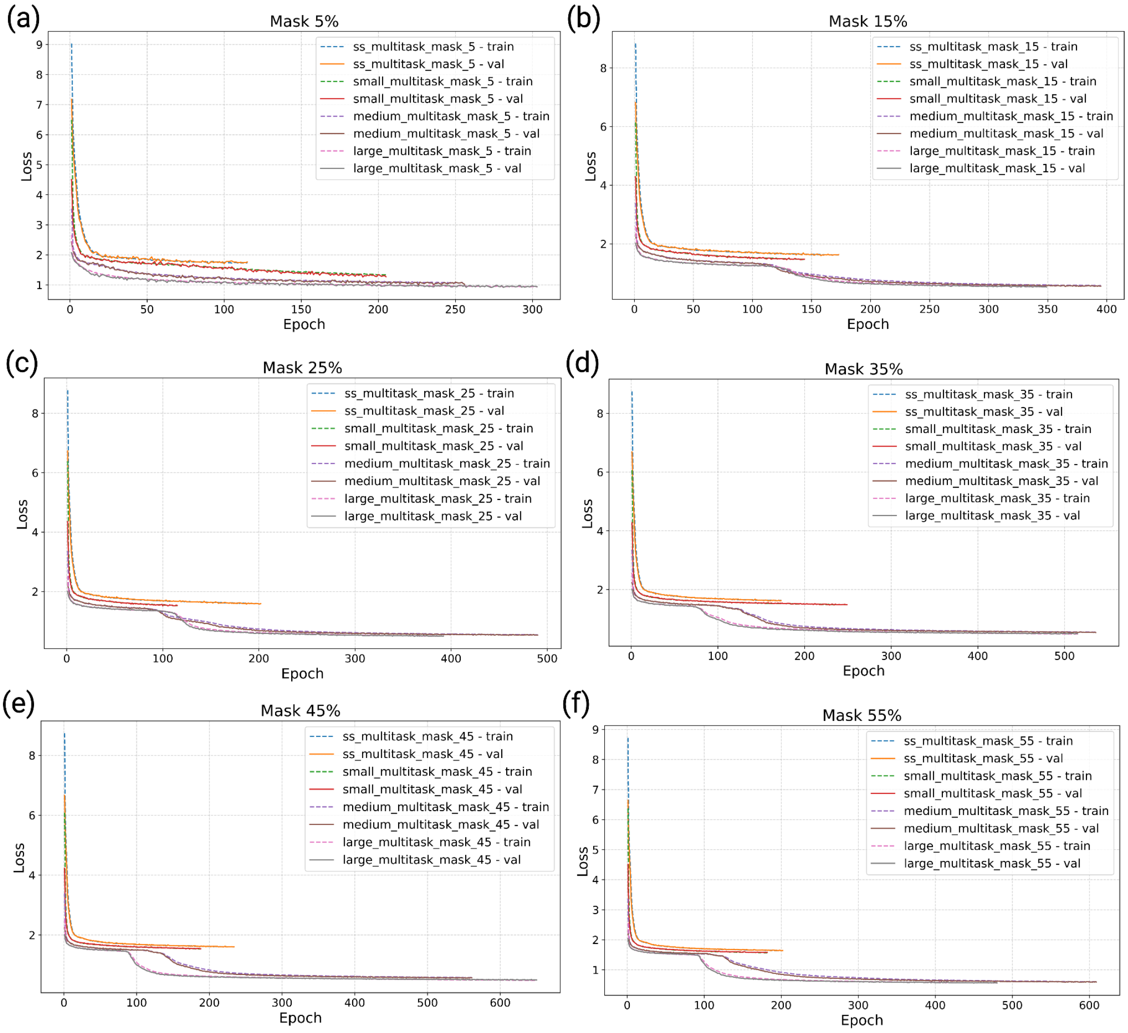


Training and validation loss curves at different masking ratios applied to nodes and edges: 5% (a), 15% (b), 25% (c), 35% (d), 45% (e), and 55% (f). For each masking ratio, models of four different sizes (ss, small, medium, and large) were trained.

Supplementary Figure S3: Comparison of prediction performance across masking ratios for different tasks

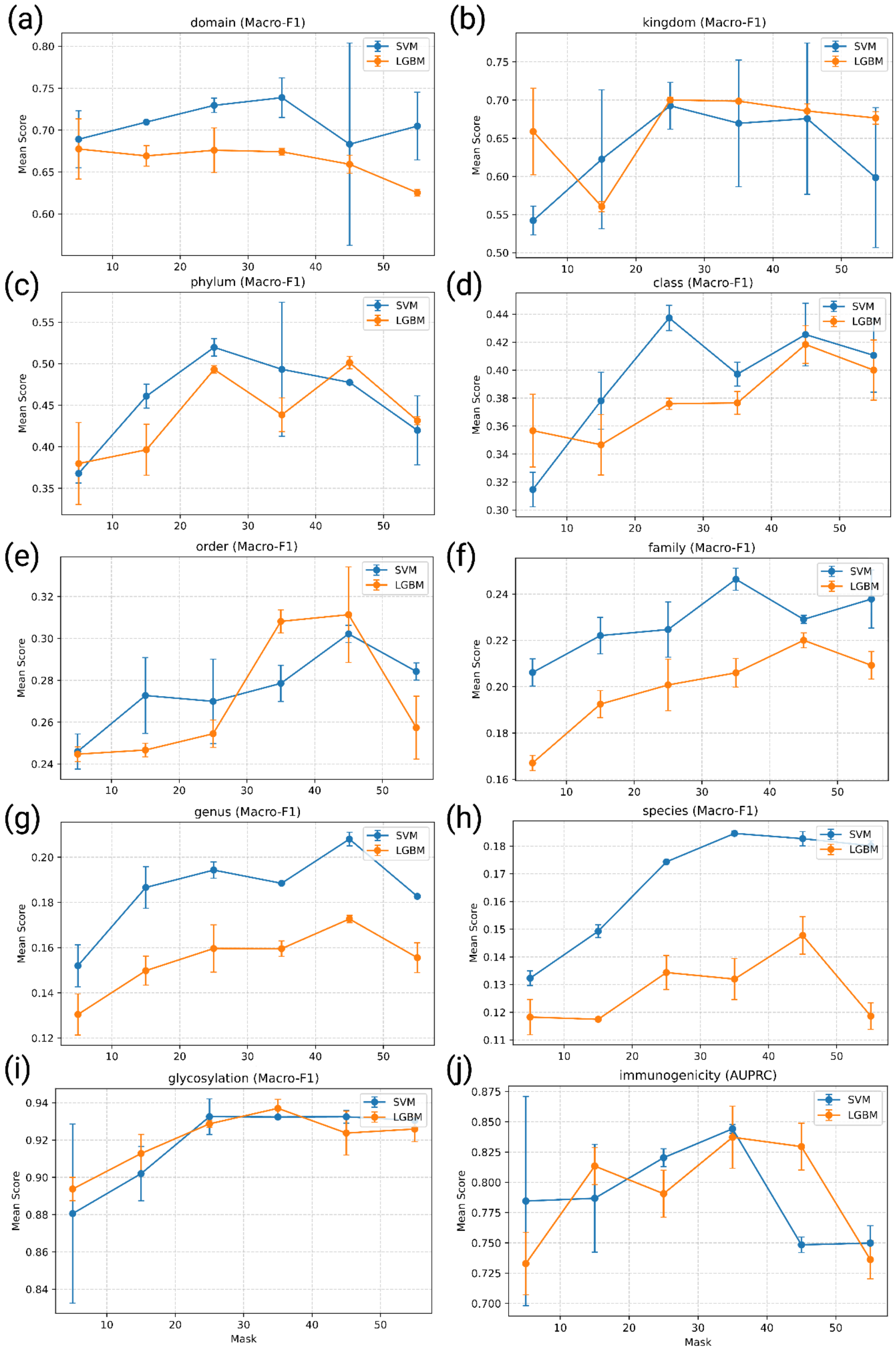

Prediction performance under different masking ratios. (a–h) Results for taxonomy classification tasks, (i) glycosylation prediction, and (j) immunogenicity prediction. Metrics used were Macro-F1 for taxonomy and glycosylation (a–i) and AUPRC for immunogenicity (j). Predictions were performed using support vector machines (SVM) and LightGBM. Error bars indicate ± standard deviation across replicates.

Supplementary Figure S4: Accuracy comparison of GlycanGT and baseline models across prediction tasks

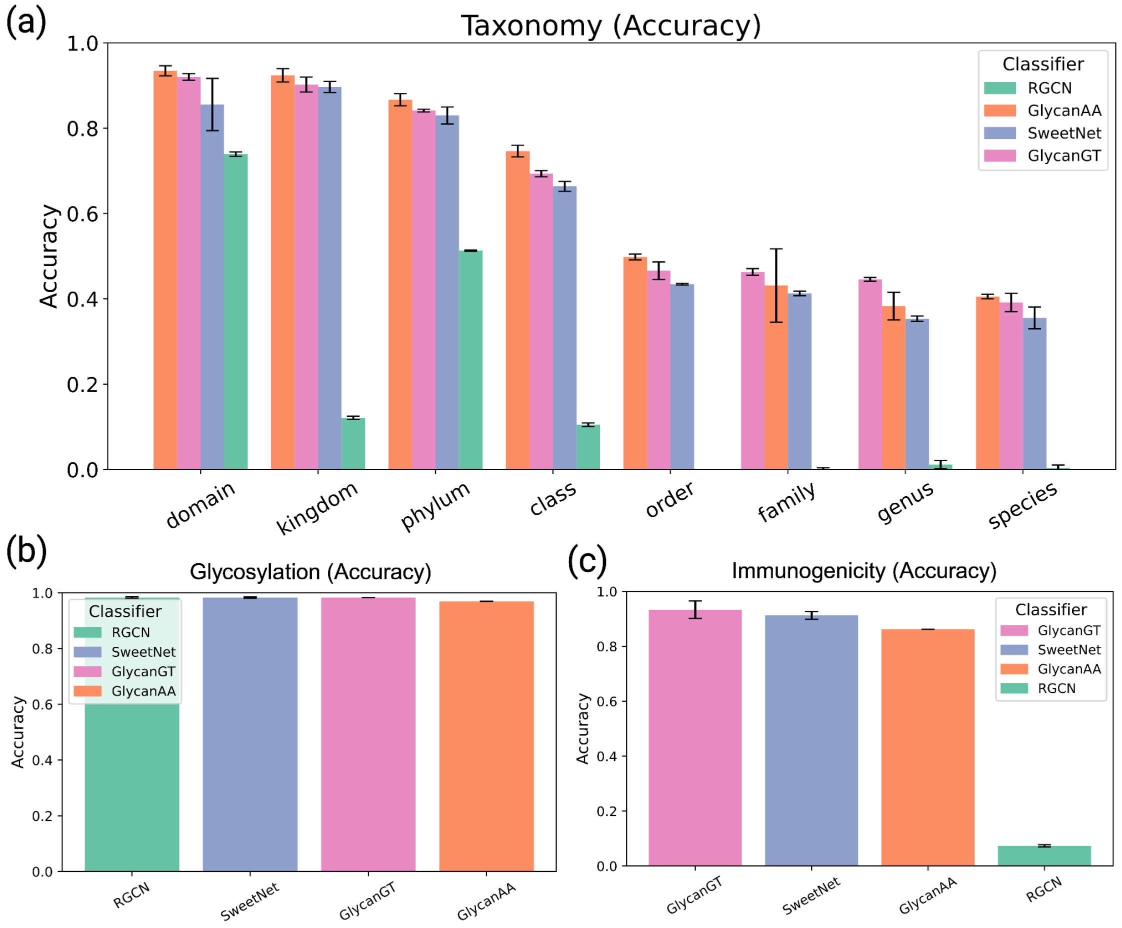


All panels were evaluated by Accuracy. (a) Taxonomy classification across eight hierarchical levels (domain to species). (b) Glycosylation prediction. (c) Immunogenicity prediction. Error bars represent mean ± s.d. across replicates.

Supplementary Figure S5: Graph kernel–based hierarchical clustering of glycans

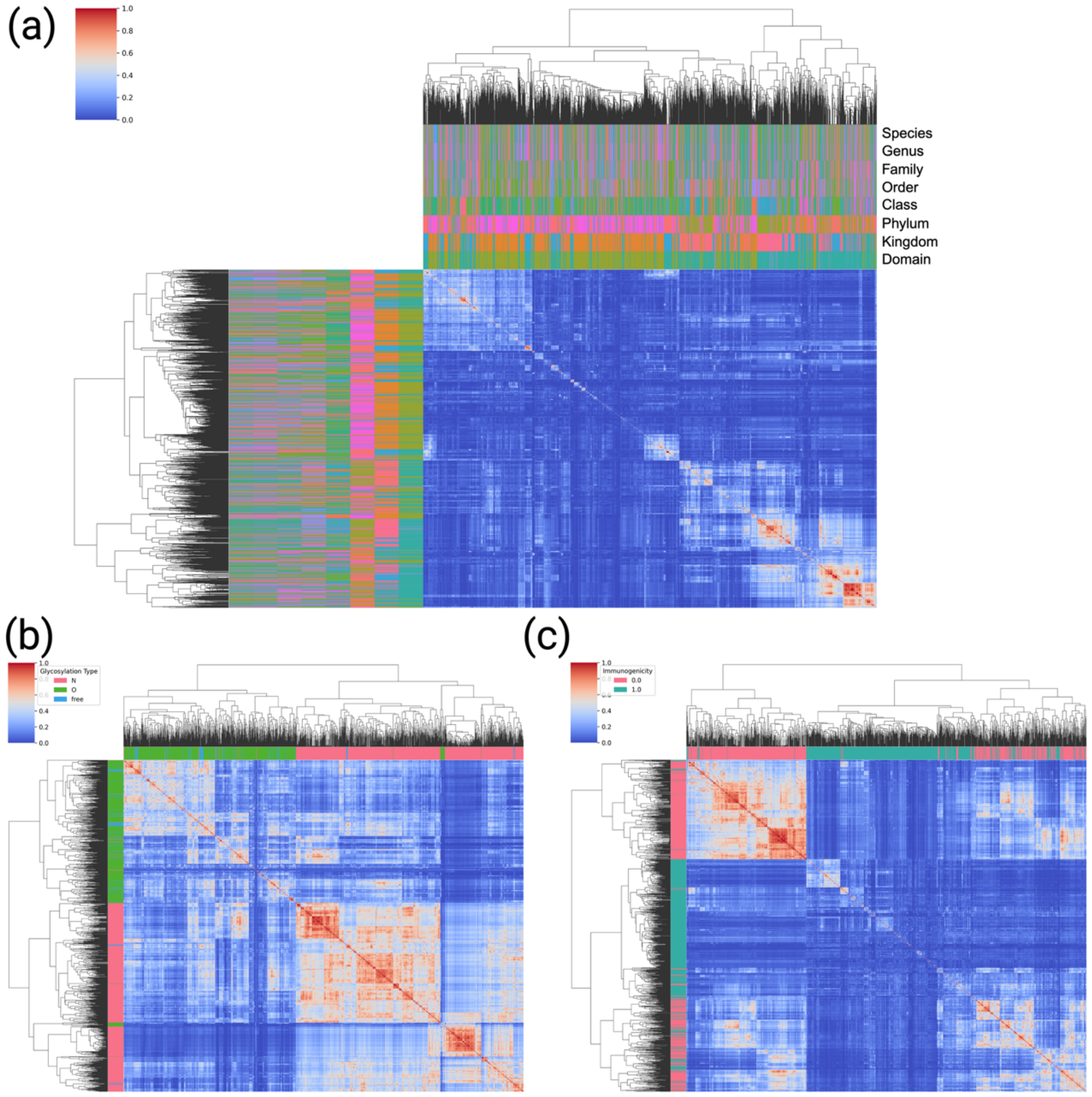


Pairwise similarities between glycans were computed using the Weisfeiler–Lehman (WL) subtree kernel and visualized by hierarchical clustering. (a) Taxonomy categories (domain to species). (b) Glycosylation types (N-linked, O-linked, free glycans). (c) Immunogenicity (immunogenic vs. non-immunogenic).

Supplementary Figure S6: Ablation study and attention visualization highlight the importance of training on both monosaccharides and glycosidic bonds

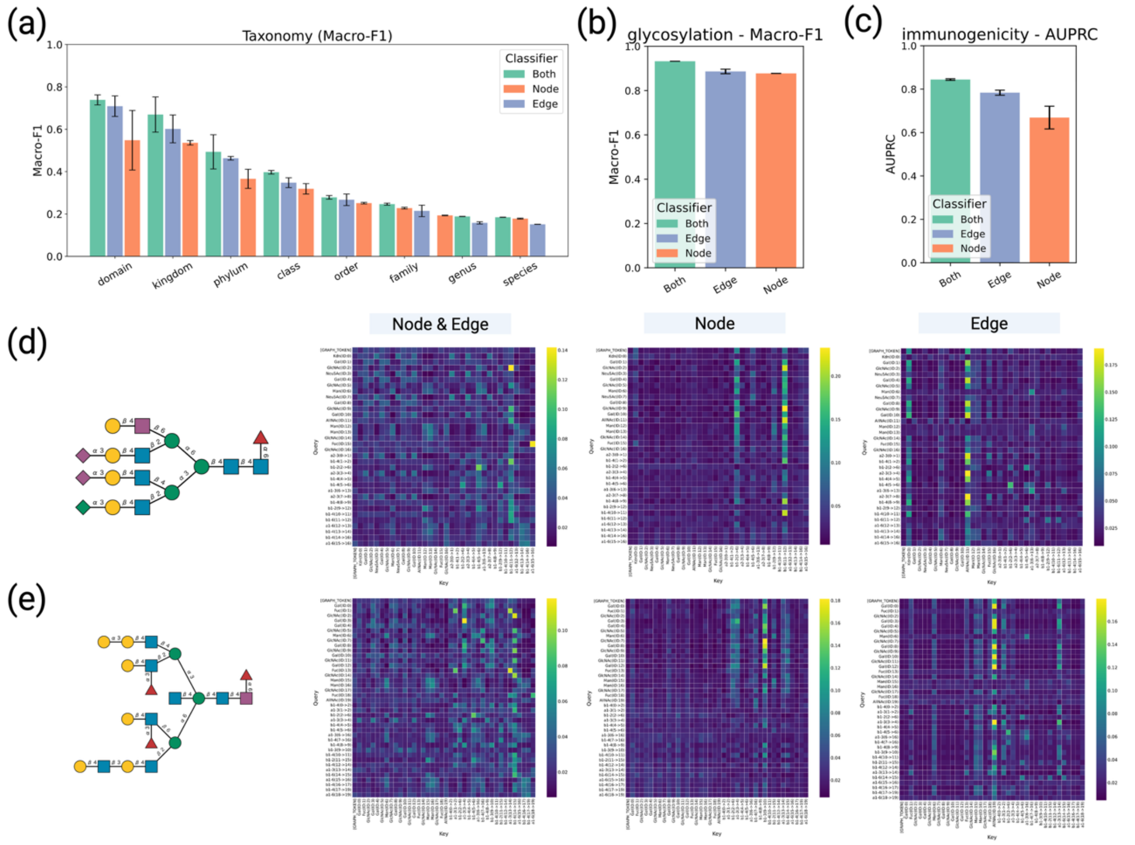


(a–c) Downstream task performance under three masking strategies: (i) balanced masking of 35% for both monosaccharides and glycosidic bonds, (ii) 70% masking of monosaccharides only, and (iii) 70% masking of glycosidic bonds only. (A) Taxonomy classification evaluated by Macro-F1 across eight hierarchical levels (domain to species). (b) Glycosylation prediction evaluated by Macro-F1. (c) Immunogenicity prediction evaluated by AUPRC. Error bars represent mean ± s.d. across replicates. (d, e) Heatmaps of attention weights illustrating token–token relationships. The left panel shows the glycan structure, followed by attention heatmaps from the same three models: (i) balanced masking of 35% for both token types, (ii) 70% monosaccharide masking only, and (iii) 70% glycosidic bond masking only. In each heatmap, the vertical axis represents query tokens and the horizontal axis represents key tokens.

Supplementary Figure S7: Distribution of glycans with ambiguous information in GlyTouCan

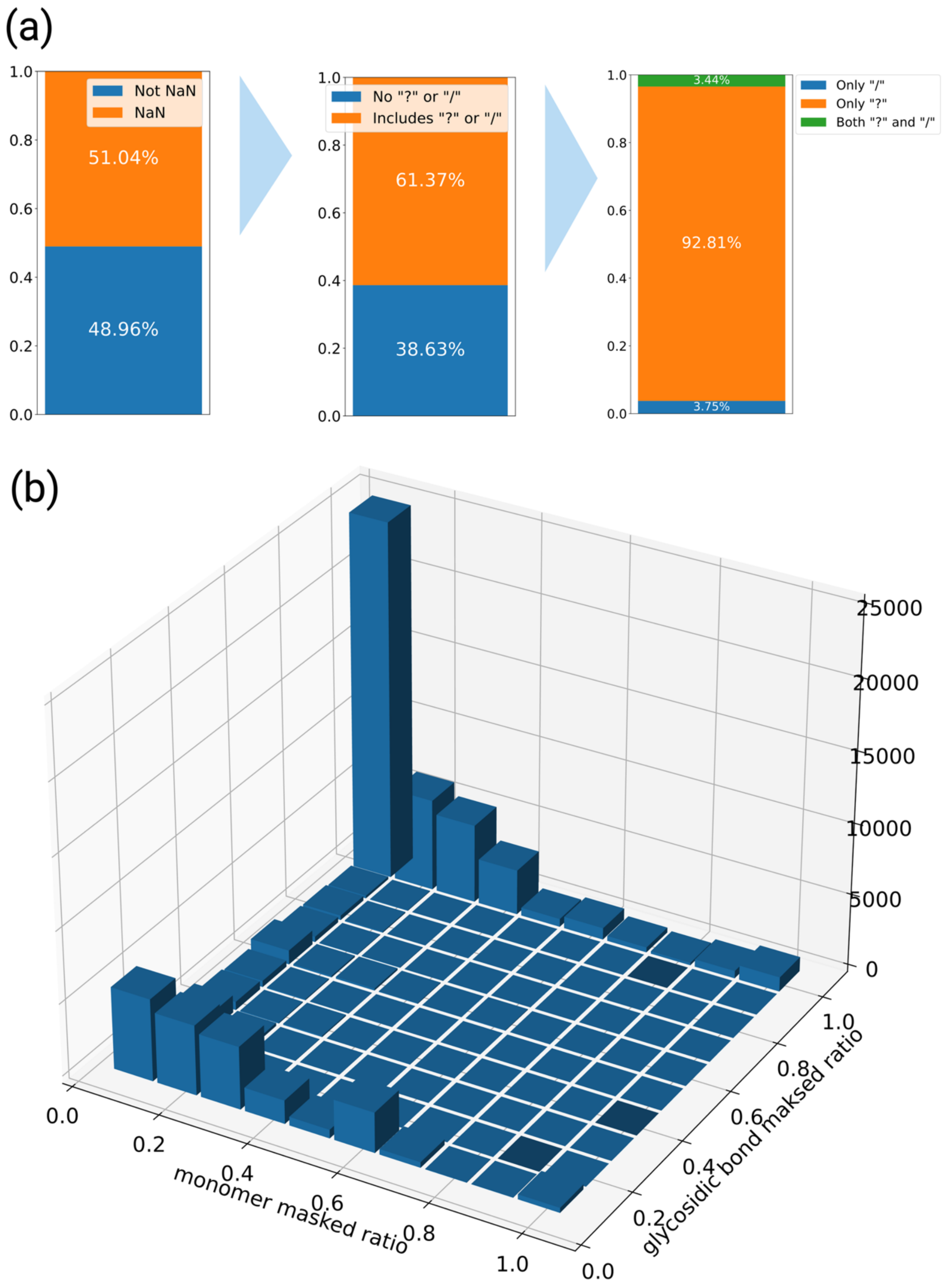


(a) Proportion of glycans in GlyTouCan (n = 244,842) containing ambiguous annotations.

Shown are the fractions of glycans with missing values (NA) and those containing ambiguous symbols such as “?” or “/”. (b) Distribution of ambiguous annotations (“?”) across monosaccharide and glycosidic bond positions among glycans containing at least one ambiguous symbol.
